# Seasonal dynamics of microbial communities mediate aroma and flavour formation during palm sap fermentation

**DOI:** 10.64898/2026.07.12.737599

**Authors:** I Nyoman Sumerta, Kate Howell

**Affiliations:** School of Agriculture, Food and Ecosystem Sciences, The University of Melbourne, Victoria 3010, Australia; Research Center for Biosystematics and Evolution, National Research and Innovation Agency (BRIN), Jakarta Pusat 10340, Indonesia

**Keywords:** spontaneous fermentation, microbial dynamics, palm wine, sap tapping, season

## Abstract

In many tropical countries, fermentation of palm sap into palm wine is an important fermented beverage contributing to local economies, tradition, and culture. Traditionally made in villages and families, palm sap is not inoculated with starter cultures and fermentation commences spontaneously. It is therefore possible that fermentation is influenced by multiple ecological factors, which affect microbial dynamics and thus flavour outcomes. Here, we studied microbial communities during fermentation of palm sap from three different palm tree species (palmyra, coconut, and sugar palm) on the island of Bali, Indonesia in both the wet and dry seasons. Our results suggest that season of collection has a strong influence on microbial dynamics and succession, and these changes positively correlate to metabolite concentration. The change of the season from the dry to wet season led to the loss of microbial diversity with lower richness in the dry season. The dominance of *Saccharomyces cerevisiae* was not affected by season and fermentation time and was dominant in all samples. Potential spoilage species, such as *Candida tropicalis* were negatively correlated to ester production and more abundant in the dry season. As microbial species varied in incidence and thus biochemical activity, the chemical groups of esters from their metabolism related to the change of season and fermentation time, while volatile compounds and small molecules were highly discriminated by season in the resultant wines. Ethyl octanoate was consistently different across all variables through comparison by three-way ANOVA and is proposed as a biomarker of seasonal variation in palm sap fermentation. These findings improve our understanding of microbial dynamics in palm sap fermentation, revealing flavour differentiation within season and suggests that strategies for microbial management, product development and quality assurance will elevate this traditional product into the future.

## 1. Introduction

Microbial communities play decisive roles in the conversion of plant substrates into food for human cultures around the world. The range of microbial interactions within a food ecosystem demonstrates microbial adaptation and competition within these dynamic substrates. Microbes, especially yeasts and bacteria, exist in food environments using specialised metabolic strategies to dominate the microbial communities in fermented food and beverages, which may be constructed and constrained by the metabolic processes (D. Liu et al., 2021; Pérez-Torrado et al., 2017; Silverstein et al., 2024). Microbes often demonstrate spatial and temporal dynamics where the complexity of food substrate and the surrounding environment affects microbial species occurrence (Barata et al., 2012; Conacher et al., 2021). Numerically dominant species are often deployed as starter cultures to conduct a fermentation and often co-exist with other species to generate flavour advancement (Steensels et al., 2019), but the use of starter cultures overrides many of the microbial dynamics that take place.

Ecological determinates affect the regional characteristics in grape wine products (Bokulich et al., 2016; Knight et al., 2015). From a microbial perspective, ecological characteristics such as raw material composition and temperature can alter the fermentation process, resulting in products with distinctive aromas and flavours while regional ecological characteristics (such as sunlight interception, soil, aspect, plant species, microclimate) create a product with a unique sensory experience and propose an economic value (Bokulich et al., 2016; Fiore-Donno et al., 2024). Local climatic factors such as temperature, precipitation, and rainfall across seasons can shape the composition of soil and plant associated microbes (Classen et al., 2015; Ding & Melcher, 2016; He et al., 2024; D. Liu et al., 2019; Waldrop & Firestone, 2006). This reflects that annual weather cycles or season contributes to the dynamics of microbial communities attached in plant-based food and beverage products.

Palm sap is a multipurpose palm product used for making palm wine, a traditional fermented beverage, that is often found in tropical countries across the globe. Inflorescence sap from coconut, palmyra, sugar palm, nipa, date palm trees are often used for the production of palm wine (Hebbar et al., 2018; Sumerta et al., 2025). Palm sap can be used to produce various food products, however its volume and composition are significantly affected by season (Sumerta et al., 2025; Ziadi et al., 2014). Produced primarily in the equatorial regions, palm wine is made throughout the seasons from the wet to dry season. Palm wine is generally fermented in open vessels, and the fermentation arises spontaneously by microbial communities without the addition of defined cultures, meaning the production process is influenced by many ecological factors, leading to microbial dynamics which likely affect the metabolite composition and thus flavour of palm wine. However, there are limited studies that address the direct impact of seasonal pattern to spontaneous fermentation, how season can affect microbial prevalence and diversity and the influence of different tree species. In other studies, diverse fermenting microorganisms in palm wine alter the chemical composition of the resultant palm wine, which is likely to alter flavour and aroma development (Das & Tamang, 2023; Delgado-Ospina et al., 2026; Djeni et al., 2022). However, the dynamics of these microbial ecosystems have not been well-understood and not well connected to metabolite production during an annual cycle of production.

Palm wine fermentation is dominated by the yeast *Saccharomyces cerevisiae* yet its interactions with non-*Saccharomyces* yeast also present in the fermentation are not well understood (Das & Tamang, 2023; Djeni et al., 2020, 2022; Sumerta et al., 2025). Synergistic metabolism and interactions between *Saccharomyces* and non-*Saccharomyces* occur in many fermented food and beverage, and strongly affect the organoleptic properties of the product. Ciani & Comitini (2015) report that non-*Saccharomyces* use amino acid and provide nitrogen source to *Saccharomyces* at the early stage in wine fermentation, then lactic acid and acetic acid bacteria contribute to the acidic fermentation and play significant roles in aroma and flavour distinction. In palm wine fermentation, lactic acid bacteria (LAB) are present during fermentation (Kouame et al., 2020) and the microbial diversity shows a significant impact on the final product of sap fermentation characters (Das & Tamang, 2023; Djeni et al., 2022; Sumerta et al., 2025). Yet, we do not know the fermentation characteristics across seasons that are linked to the microbial and metabolite composition affecting aroma and flavour of the final product. Thus, it is important to understand microbial diversity and abundance during sap fermentation, which will reveal the construction of aroma and flavour over different seasons.

In this study, we sampled palm sap samples from across villages in Bali, Indonesia to capture the extent of microbial diversity in traditional production processes. We collected samples in both the wet and dry seasons and from three different palm species (palmyra, coconut, sugar palm) and allowed the sap to ferment naturally with regular sampling. We performed DNA amplicon sequencing to identify microbial communities of fungi and bacteria and measured fermented palm sap metabolites at each fermentation stage. We observed connections between microbial communities and metabolites, which was altered by season of sap collection. Our results contribute to understandings of the effect of seasonal factors in spontaneous fermentation in this important traditional beverage.

## 2. Materials and Methods

### Collection and sampling of fermenting palm sap

Fresh palm sap samples were collected from overnight sap tapping process in the villages of Bali, Indonesia. Ten villages were selected to demonstrate the range of palm species and season collection, considering the accessibility, availability, and recognition of well-known villages that produce palm wine or palm spirits. There were three commonly-tapped palm species sampled in Bali: coconut (*Cocos nucifera*), palmyra (*Borassus flabellifer*), and sugar palm (*Arenga pinnata*). We worked with farmers to collect the sap. The farmer would climb the palm tree and collect sap from a cut inflorescence that had been collecting sap for approximately 24 hours. When the farmer returned to the ground, we collected the sap. Here, we took 300-500 ml of fresh palm sap, coarse filtered with a sterilised strainer to remove unwanted particulate matter before storing in a sterilise plastic container on ice before returning to the laboratory. Once in the laboratory, the samples were returned to room temperature and left to naturally ferment for a week at room temperature (between 27 and 30°C). For sampling, approximately 10 mL of the fresh palm sap was collected into a sterile centrifuge tubes, at the start of fermentation (after the first day; 1D), on the third (3D) and seventh day (7D) after collection. From an individual village, three separate samples were taken for fermentation in the laboratory, with single samples taken at each time point. All collected samples were frozen for microbial and metabolic analyses.

To obtain the influence of seasonal tapping process, sample collection took place in both the wet and dry seasons with similar collection procedures. The first collection was between late October and early November 2022 (early wet season) and the second batch was in July 2023 (middle of the dry season). These collection times were based on compiled weather data (**Supplementary Fig. S1**), showing rainfall rate and sunlight exposure were high during July-August for dry season collection, while rainfall during October-November started to decline for wet season collection. The early wet season was selected due to the difficulty in climbing trees to collect the sap during heavy rain experienced later in the wet season. In total, 27 samples of fresh palm sap were gathered at nine different villages (triplicate samples were taken in every village) in the wet season, yielding 81 samples after the fermentation. The same number was gathered for the dry season, but with one different village included (Lebu; L) and one village was excluded (Gempinis; GM) due to accessibility issues and availability of palm sap sample during the dry season. Details of collection sites are available in **Supplementary Table S1**.

### DNA extraction and amplicon sequencing

Extraction process of DNA was performed with pre-treatment and using commercial extraction kit (PowerSoil, Qiagen) as described by Sumerta et al. (2024). Briefly, 1 mL of sap sample was centrifuged to obtain pellets of microbial cells, then washed twice with polyvinylpolypyrrolidone (PVPP) in 0.1 M phosphate-buffered saline (pH 5.2) before following the extraction kit procedures. Gel electrophoresis was used to clarify gDNA quality and Nanodrop spectrophotometer and Qubit (Thermo-Fisher Qubit 2.0 Fluorometer) for quantitatively measuring DNA concentration. If DNA in a sample was found to be poor quality, we further treated the sample by precipitating DNA with absolute ethanol and 3 M sodium acetate, centrifuging and then resuspending in 100 μL volume with TE buffer. All samples were then stored at –20°C for amplicon sequencing at 16S rRNA hypervariable region (V3-V4; 341F – 5’ CCTACGGGNGGCWGCAG 3’, 805R – 5’ GACTACHVGGGTATCTAATCC 3’) for bacterial communities and ITS1 region (ITS1F – 5’ CTTGGTCATTTAGAGGAAGTAA 3’, ITS2R – 5’ GCTGCGTTCTTCATCGATGC 3’) for fungal communities using the standard service from the Australian Genome Research Facility (Victoria, Australia).

### Metabolite profiling

The general characteristics of palm sap samples were measured followed by metabolic profiling technologies to examine volatile organic compounds and both volatile and non-volatile small molecules in the palm wine.

#### a. Basic chemical parameters

FTIR-based analysis using an OenoFoss^™^ wine analyser (FOSS Analytical co., Denmark) was used to measure chemical compounds related to total sugar content, titratable acidity (TA), and pH, calculated using the finished wine calibration.

#### b. Volatile organic compounds by HS-SPME-GC/MS

Samples for volatile analysis were prepared according to Luo et al. (2022) with some modifications for Head Space-Microextraction–Gas Chromatography and Mass Spectroscopy (HS-SPME-GC/MS). Here, approximately 5 ml of each sample were prepared in 10 ml HS-GC vial with magnetic cap, containing 1 g of NaCl to increase the volatility and then 100 µL of 4-octanol (2.5 mg/L) were added as internal standard (IS). Incubation was performed to sample vial in agitator for 15 minutes at 40°C prior fibre extraction. A 65 μm PDMS/DVB, 10 mm fused silica fibre was exposed to vial with agitation for 30 minutes. The fibre was then desorbed in splitless mode to GC-MS (Agilent 8890GC-59778B MSD equipped by autosampler PAL RSI 85), fitted in a DB5 column (30 m x 0.25 m x 0.25 µm; Agilent J&W) with helium as a carrier gas and the flow rate maintained at 1 mL/minute. GC-MS was configured according to the parameters given by Borse et al. (2007) through setting the injector port temperature for fibre desorption at 220°C. Oven temperature was started at 35°C for 2 minutes and then increased to 90°C at the rate of 1.5°C/minute and further increased to 220°C at the rate of 5°C/minute. Data acquisition was configured through MS Source transfer line at 230°C and 150°C for MS Quad with scanning mass acquisition mode in range of 35-350 m/z. For absolute quantification, serial external standards (ES) were analysed by incorporating to mock palm wine (5% v/v ethanol, 0.5% w/v acetic acid, 1% w/v gallic acid, 0.5% w/v Na=SO=, and 1% w/v citric acid) and these peak areas were used for constructing calibration curves.

#### c. Volatile and non-volatile small molecules by ^1^H NMR

Proton nuclear magnetic resonance (^1^H-NMR) was used for a wider small molecules as described by Winters et al. (2022). Briefly, samples were centrifuged at 10,000 x *g* for 1 minute and the supernatant was taken for analysis. A 0.15 M final concentration of phosphate buffer was used as a solvent, containing sodium phosphate monobasic (NaH_2_PO_3_), sodium phosphate dibasic dihydrate (Na_2_HPO_3_, 2 H_2_O), sodium azide (NaN3), and 3-trimethyl-silyl-[2,2,3,3-2H4] propionic acid sodium salt (TSP-d4, 98 atom % D) diluted with MilliQ water, then filtered using 0.22 Millipore filter. About 800 µL of phosphate buffer were mixed with 200 µL of D_2_O (8:2). The mixed buffer was transferred about 300 µL to 700 µL of sample (3:7) into a new tube and vortexed vigorously before 700 µL of the mixture was added to 5 mm O.D. NMR SampleJet tubes (Bruker, Germany). NMR spectra were measured on a Bruker Avance NEO system at 500.13 MHz proton frequency. The machine was equipped with TXI cryoprobe for greater proton sensitivity, an autosampler and a broadband probe. Data acquisition was set to the relaxation delay (d1) for 2 seconds. The standard pulse sequence was used for measuring NMR spectra at 298 K with zgcppr pulse program, sweep width at 7812.500 Hz, and acquisition time for 2 seconds. Automatic phase and baseline processing were performed in the TopSpin software version 4.2 (Bruker, Germany).

### Data and statistical analyses

*Basic chemical analysis*: chemical data was exported from FTIR Oenofoss machine into.xls file, manually summarised as briefly displayed in range value in **Supplementary Table S2**. The FTIR metadata was then imported to R ver. 4.4.0 in RStudio 2025. 09.02+418 with ggplot2, tidyverse, and reshape2 packages for bar chart visualisation.

*Amplicon sequencing data analysis:* sequenced amplicons were quality checked by FastQC before and after adapter trimming using BBDuk in Geneious with parameters in trimming both ends, minimum quality at 30, *K*mer length at 27, and minimum length at 80 bp. All paired-end trimmed sequences were demultiplexed and denoised in Qiime2 (Bolyen et al., 2019) through DADA2 to infer non-chimeric amplicon sequence variants (ASVs) (Callahan et al., 2016), subsequently aligned with Naive Bayes feature classifier of Greengenes 2022.10 database for bacterial ASVs (McDonald et al., 2024) and UNITE 2024 database for fungal ASVs (Abarenkov et al., 2024). The Qiime2 results were then constructed to phyloseq object (McMurdie & Holmes, 2013) in RStudio with qiime2R, phyloseq, ggplot2, DESeq2, microbiome, tidyverse, reshape2, vegan, ggpubr, and microbiomeutilities packages for data visualisation and statistical analysis. To maintain data reproducibility and to obtain a more balanced comparison and composition of ASVs, sampling depth was taken for bacterial (32,861) and fungal amplicons (6,549) based on the sample feature characteristics. Statistical pairwise comparison was performed through Wilcoxon test with false discovery rate (FDR) *p*adj = 0.05, ANOSIM test for multivariate comparison, and DESeq2 for differential analysis (Love et al., 2014).

*HS-SPME-GC-MS data analysis:* to construct the representative compound pattern for autointegration, about 10% of total samples representing palm species, season, and fermentation stage were overlayed and integrated with RTE parameters for area counts > 24,000 to equivalent with IS peak area. Compound candidates were aligned based on retention time and selected according to mass spectra similarity (>70%) from the National Institute of Standards and Technology library. Compounds consistently presented in blanks were removed from further analysis. Individual selected compounds were compared to retention indices (RI) after mixed alkane standards (C6-30) were analysed for supporting compound identification (score difference >110, discarded). The configured autointegration was then applied to all samples. Compounds that were detected <20% and <10,000 mean peak abundance with median = 0 of total samples were discarded. Further identification and absolute quantification were achieved by comparing the retention time and mass spectra similarity of the external standards. Quantification was done by calculating ratio of compound and IS peak areas before being subjected to the slope of standard calibration curves. For compounds not corresponding to the external standards, semi-quantification was addressed to peak ratio equivalent to IS concentration (μg/L). Data was then constructed to multivariate, discriminant, and correlation analyses in RStudio using ggplot2, tidyverse, reshape2, ggpubr, and factoextra packages.

*^1^*H*’NMR spectral data analysis:* spectral processing and analysis were performed in NMRProcFlow software (Jacob et al., 2017). The spectra processing was performed following software metabolite fingerprinting procedures and ppm shift was referenced to TSP-d4 (δ=0.00). The spectral baseline correction was adjusted by global and local correction on soft level. The alignment step of the spectra was performed with Least-Squares algorithm comparing average spectrum as reference and then realigned with Parametric Time Wrapping (PTW) to deal with ambiguous peak alignment. The alignment spectra binning process was done by intelligent bucketing method, splitting area with the same resonance with SNR threshold at 3 (De Meyer et al., 2008). Data matrix was then normalised with TSP ppm range and subsequently loaded to Biostatflow v.2.9.6 (https://biostatflow.org/) for scaling, hierarchical clustering analysis (HCA), correlation, discriminant modelling, and multivariate analysis. Clustering data was then revisualised in RStudio for PLS-DA plots (using packages mdatools and ggplot2). The identification of the compounds was based on a list of ppm obtained from clustering and multivariate analysis. The list was then compared to metabolite databases in NMR PeakMatching 1.0 platform (https://pmb-bordeaux.fr/PM/webapp) and PeakForest Alpha (https://alpha.peakforest.org/), and the matching results are available in **Supplementary Table S3**.

## 3. Results

### 3.1 Seasonal shifts in palm sap chemical composition

We obtained 162 samples of palm wine, representing two seasons of sap collections, three fermentation time, three palm species in ten villages. Freshly collected, palm sap is slightly viscous, translucent, and sweet. Bees, wasps, and flies were often trapped and floating in the collection vessel and were removed with a sieve for palm wine fermentation. In some areas, the production of palm wine was enriched by a natural booster, such as coconut husks and pieces of specific woods to improve alcohol production and organoleptic properties. These were removed before sap fermentation. Sampling sites, sap tapping process, fermentation, and fresh sap general chemical profiles are presented in **Fig. 1**. Sampling locations were distributed in villages located in the eastern part of the island of Bali (**Fig. 1C**), where production of palm wine is very well-known in local communities. The sap was collected by climbing the tree and cutting the tip of palm blossom/flower, where the sap collected in a container overnight (**Fig. 1B**). In the morning, the fresh sap is generally collected and usually left to ferment or sold fresh by the farmers.

**Fig. 1.**
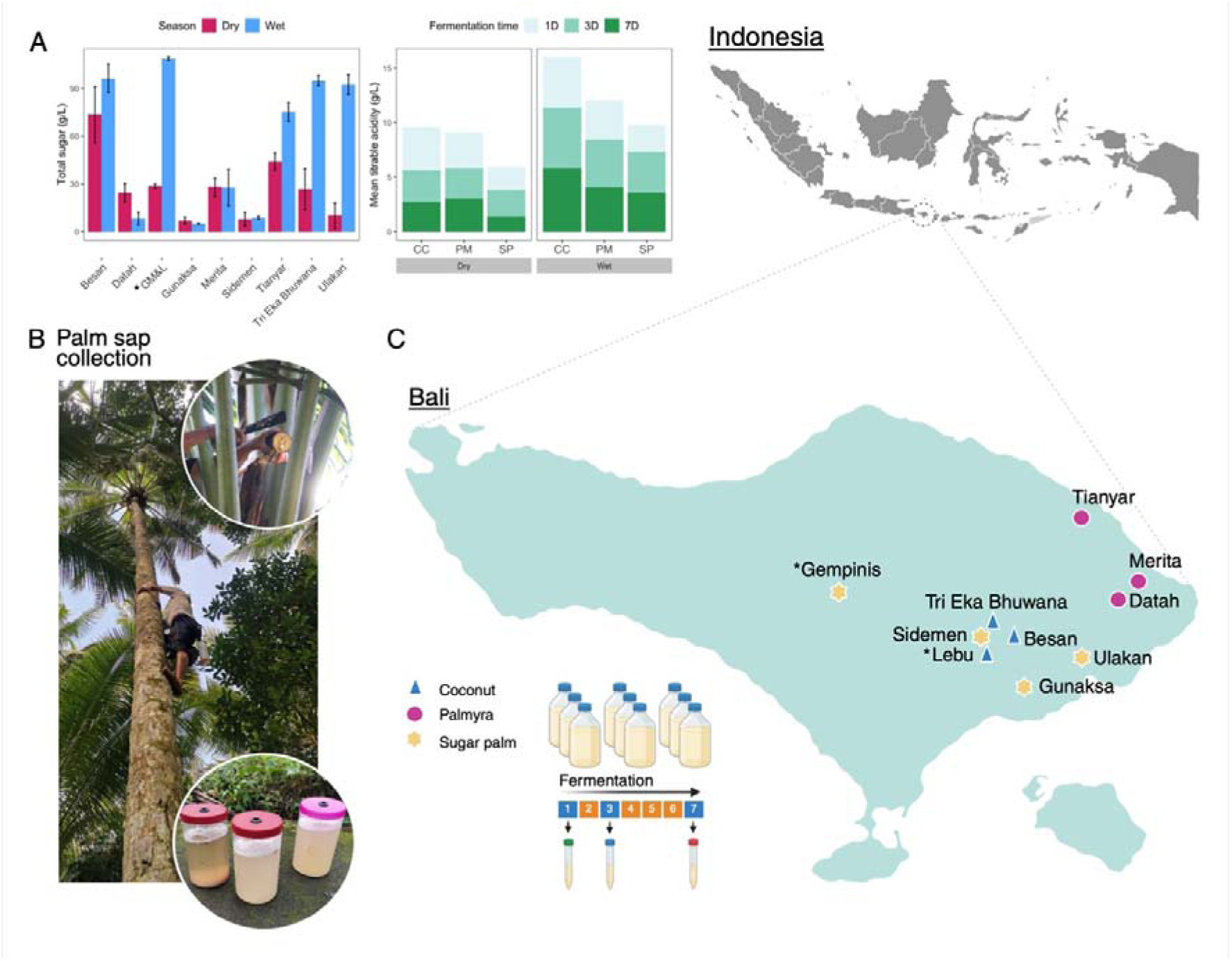
Palm sap collection, sampling sites and palm sap chemical profiles. A) Total sugar amount of fresh palm sap and titratable acidity across fermentation stages, season, and palm species. Gempinis and Lebu villages were combined (GM&L) due to partial sampling on each season. B) Farmer climbing a palm tree to cut the palm blossom, then to collect fresh palm sap. C) Map indicating the villages and palm species in sampled Bali, Indonesia with fermentation sampling strategy. A (*) indicate the partial sampling in village for each season, detail village collection in **Supplementary Table S1**.

We used FTIR-based analysis to get a general chemical profile of fresh and fermented palm sap, providing a prediction of total sugar, ethanol, and organic acids. However, ethanol did not reach the calibration range of our method (<10% v/v) and so was not included in further analysis. The amount of total sugar and titratable acidity (TA) varied across the collection being higher in the wet season and differing between palm species, and day of fermentation (**Fig. 1A**). Between 19.7-49.5 g/L of total sugar were detected in palmyra sap (PM), glucose 10.2-21.3 g/L, TA 0.7-12.7 g/L, and pH in range of 3.52-4.58 in the dry season, while in the wet season, total sugar was in range 8.5-81.5 g/L, glucose in 7-21.3 g/L, 0.6-5.2 g/L, and pH 3.53-4.04. Sugar palm sap (SP) had total sugar about 3.8-18.1 g/L, glucose 2-14.1 g/L, titratable acidity 0.9-3.2 g/L, and pH in range 3.81-4.24 in the dry season, while in the wet season, total sugar was in the range of 4.7 to 109.1 g/L, glucose 2.1-7.9 g/L, TA 1.1-4.3 g/L, and pH 3.17-3.91. Coconut sap (CC) had between 13.7 and 91.1 g/l of total sugar, glucose was between 1.8 and 22.2 g/L, while TA was 1.3-14.6 g/l, and pH 2.3-4.06 in the dry season. In the wet season, coconut palm sap had total sugar between 85.7-101.6 g/L, TA 1.2-7.8 g/L, and pH 3.81-4.13. Details of chemical data from FTIR including fermentation stage are presented in **Supplementary Table S2**. The chemical profile of fresh palm sap was significantly impacted by the change of season, reflected in the amount of total sugar and TA in most of the collection villages and fermentation time.

### 3.2 Microbial community structures vary during palm sap fermentation

Amplicon sequencing provided a targeted microbial profile from two diverse regions of bacterial and fungal conserved genes. A total of 23,805,748 high quality bacterial V3-V4 region sequences were obtained after pretreatment in adapter and primer trimming, denoising, and chimeric detection of 162 samples. Number of amplicon sequence variants (ASVs) for bacteria were clustered to 7,082 features, classified to two predominant phyla of Proteobacteria and Firmicutes. The bacterial composition was depicted in the mean relative abundant (MRA), taxonomic profiles, and diversity indices, showing a large variability of genera domination and unique among all samples (**Fig. 2A**). *Zymomonas*, *Liquorilactobacilus, Acetobacter*, and *Lactobacillus* dominated the bacterial MRA. Fungal ITS1 amplification gave 16,848,329 reads that clustered to 1,990 features to Ascomycota with *Saccharomyces* and *Candida* the most prevalent genera. Amongst the data, some genera demonstrated specific dominance in certain samples and villages. For example, *Candida* was observed higher prevalence in coconut sap from Besan village (Fig 2A: red in colour of fungal communities) and sugar palm from Gempinis and Ulakan villages, while *Starmerella* (Fig 2A: green in colour of fungal communities) was found higher in abundance in palmyra sap. *Zymobacter* (Fig 2A: light green in colour of bacterial communities) was found to be prominent in the dry season collection from Tianyar village.

**Fig. 2.**
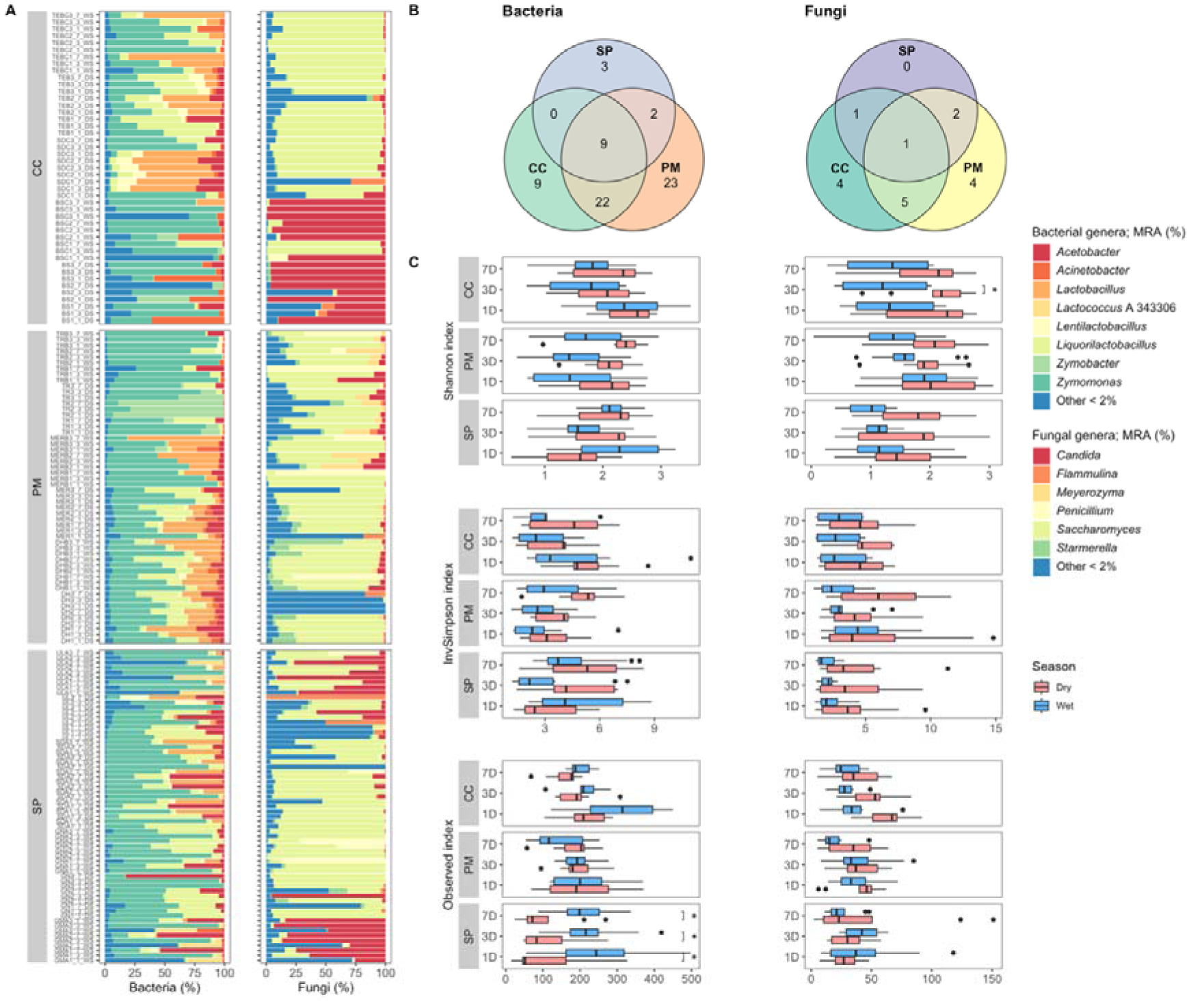
Microbial composition and diversity value of fermented sap over different seasons (dry and wet), palm species (CC, PM, SP), and fermentation time (1, 3, 7 days). A) Mean relative abundance shows species dominance (MRA >2%) in bacterial and fungal communities (Details of sample codes in Y axis: every top nine lines indicating the wet season followed by the dry season and every top 3 lines indicating a sequence of fermentation time from top day 7, 3, to day 1; CC: every top 18 lines indicating a village of Tri Eka Bhuwana (TEB), Lebu (SDC), and Besan (BS); PM: every 18 lines indicating a village of Tianyar (TR), Merita (MER), and Datah (DH); SP: every 18 lines indicating a village of Ulakan (UL), Sidemen (SDA), Gunaksa (GN), and Gempinis (GMA)) – available **on Supplementary Fig. S2**; B) Shared of the core microbial communities in each palm species; C) Alpha diversity indices show microbial diversity reduction over season (Shannon, invSimpson, Observed) but enrich the richness index.

The core and unique microbes across fermentation time and season in palm species were considered to understand the microbes were dominating the fermentation process in each palm species. The abundance and prevalence of ASVs was used to define the core microbial species with a parameter of 50% prevalence across all samples and at least 0.1% counts in each sample (**Fig. 2B**). There were nine reads of shared core bacterial ASVs across palm species, representing *Acetobacter, Liquorlactobacilllus, Zymomonas*, and *Lentilactobacillus* with palmyra showing the highest unique ASVs. Fewer unique fungal ASVs were observed among palm species due to the dominance of the core species, *S. cerevisiae,* during fermentation of all palm saps, across all sampling sites and in both seasons.

The season had an impact on microbial diversity, showing a change in diversity indices from the dry to wet season (**Fig. 2C**). Despite Wilcoxon test with False Discovery Rate (FDR)=0.01 confirming several significant pairs, the views were generally clear to the alteration of season. Only Shannon and Simpson bacteria indices in 1D of sugar palm showed higher in the wet season and also showed a significant ASVs richness in Observed index. The ASVs reads showed bacterial and fungal communities in sugar palm were higher in the wet season but unequally distributed to all samples as seen in lower value in Shannon and Simpson indices. This richness was also demonstrated over the collection villages where Observed index was significantly different, notably between the sugar palm and coconut samples in the wet season (**Supplementary Fig. S2**).

### 3.3 The influence of seasonal sap collection on microbial communities

To understand microbial composition across all samples and variables, we performed non-parametric multidimensional scaling (NMDS) analysis with Bray-Curtis’s distance to visualise community variations (**Fig. 3A-D**). Population was slightly clustered from the dense variables in NMDS plot, indicating the effects of season and palm species. In fungal communities, fermentation time contributed less to the construction of the microbial community due to dominance of core species throughout the fermentation process. These results were confirmed by the permutation analysis of similarity results (ANOSIM) showing statistical significance in each variable. Season has a significant effect on microbial beta diversity with *p*-value = 0.001 for both bacterial and fungal communities. However, the variation was limited in bacteria at 9.37% (R^2^ = 0.0937) and fungi at 11.05% (R^2^ = 0.1105). Palm species displayed higher variation of the bacterial community at 22.22% (*p*-value = 0.001) and fungal community at 29.90% (*p*-value = 0.001). The fermentation stage contributed to 4.66% (*p*-value = 0.001) in bacteria, while fungi did not show significant variation in fermentation stage (R^2^ = 0.0%), confirmed by ANOSIM test (*p*-value = 0.393). Season, palm species, and fermentation stage were the significant factors to mediate the microbial composition although no significant effect of the fermentation time was observed for the fungal community.

**Fig. 3.**
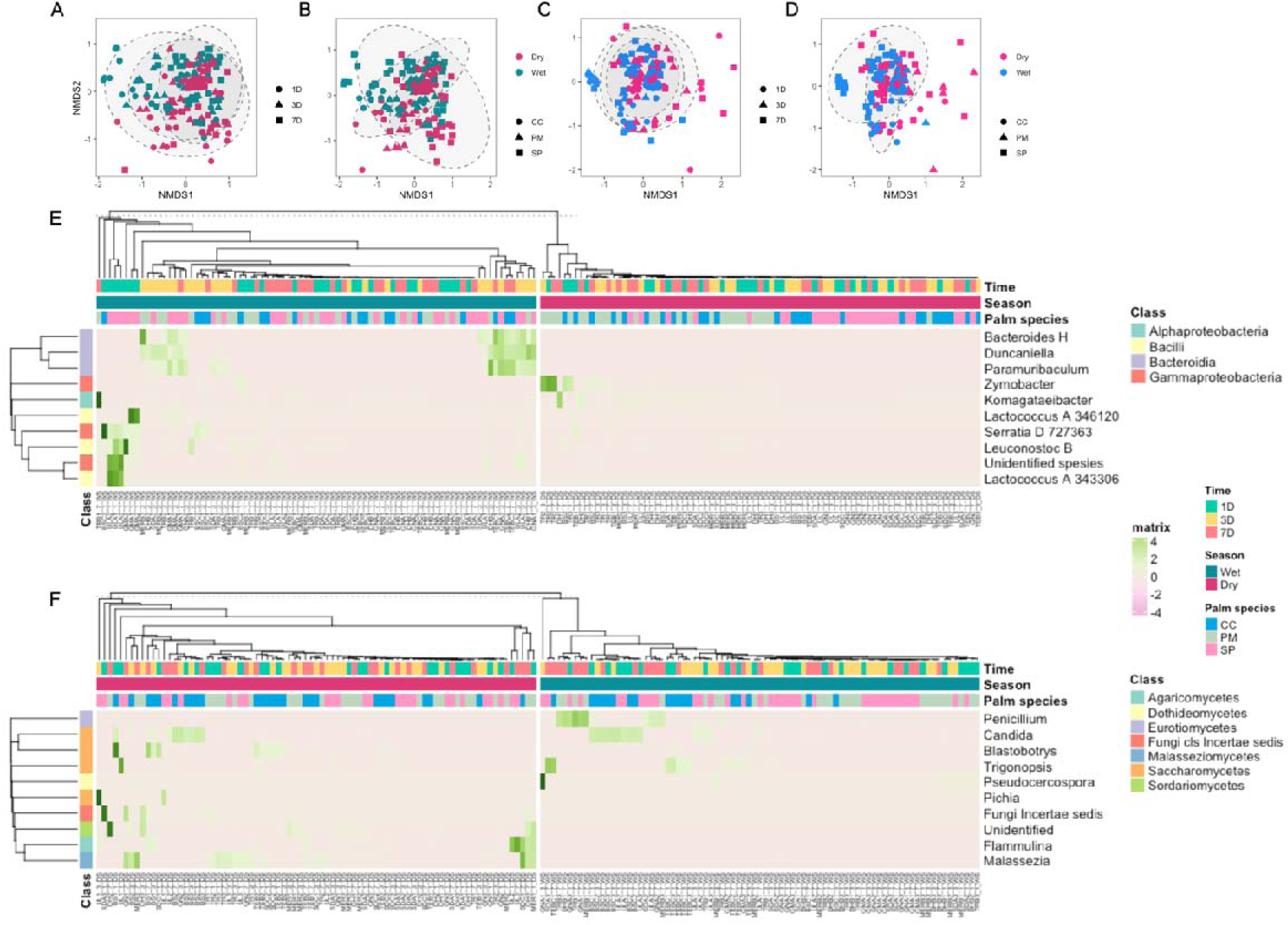
Season altered microbial communities and significantly differentiated genera composition across all studied variables (*p.adj*-value = 0.05). A) NMDS plots (ellipses 95% confidence intervals) demonstrated significant effects of season and fermentation time to bacterial composition. B) Clustered bacterial composition among palm species in different season C). Fungal composition was not separated by the fermentation time. D) Season and palm species significantly clustered fungal composition. E-F) The most significant microbial genera showing specific niche in palm species, season, and fermentation time, affecting their abundance.

We observed the alterations of season, palm species, and fermentation time can be specifically showed in genera, visualised in clustered heatmap (**Fig. 3E-F**). Through DESeq2 algorithm (*p.adj* = 0.05, log2FoldChange = 1), we selected the most significant taxa with the lowest *p.adj* value accounting 10 ASVs for bacterial genera and 10 ASVs for fungal genera. This simulation suggests that the change of dry and wet season generally differentiated the minor taxa. Lactic acid bacteria (LAB) were significantly affected by season, where *Lactococcus* and *Leuconostoc* were higher in wet season and mostly presented in fresh palm sap of sugar palm. *Bacteroides* H*, Duncaniella* and *Paramuribaculum* spp. were observed particularly in wet season of the third and seventh day of the fermentation. *Zymobacter* spp. occurred in the dry season of palmyra sap and *Komagataeibacter* spp. was also found in the dry season but significantly higher in the seventh day of the wet season in Tianyar village. In fungal communities, reads of the human-associated fungus, *Malassezia* spp., which was significantly abundant in the dry season and increased in the third and seventh day of the fermentation as was the genus *Flammulina.* In the wet season, fungi of the genera *Pseudocercospora* and *Penicillium* were significantly higher in palmyra and sugar palm sap. *Candida* spp. was apparently unaffected by season but more abundant in coconut and palmyra across the fermentation time.

### 3.4 Change in microbial composition correlates to the presence of microbial metabolites and small molecules

We analysed the volatile organic compounds (VOCs) and small molecules to view compound formation and intensity during the fermentation process. Esters were the dominant chemical group measured, making up 71.05% of total types of VOCs, followed by alcohols and benzenoids. In the PCA biplot analysis, groups of compounds were clustered by season of collection and fermentation time variables (**Fig 4A**). Ethyl decanoate, ethyl hexanoate, and ethyl octanoate were contributed to the differences between dry and wet seasons, while isoamyl ethanoate, phenylethyl acetate, and isobutyl acetate were highly correlated to the fermentation time. Decanoic acid was negatively correlated to the fermentation time. A clear cluster of VOCs was evident by season and the fermentation time, while palm species shared a similar pattern in principal component analysis (PCA) (**Fig. 4 B-C**). About 38% of the variability shared in PC1 suggesting samples differentiated in season cluster and the fermentation time. Adonis2 test confirmed that season was statistically different for differentiating VOCs with F = 0.01, R^2^ = 0.129, indicating variation of 12% across all samples. The fermentation time contributed to 13.46% variation and highly significant factor (F = 0.001), while palm species has less variation and statistical different (R^2^ = 0.042, F = 0.019), meaning less unique volatile compounds were generated by the species of palm.

**Fig. 4.**
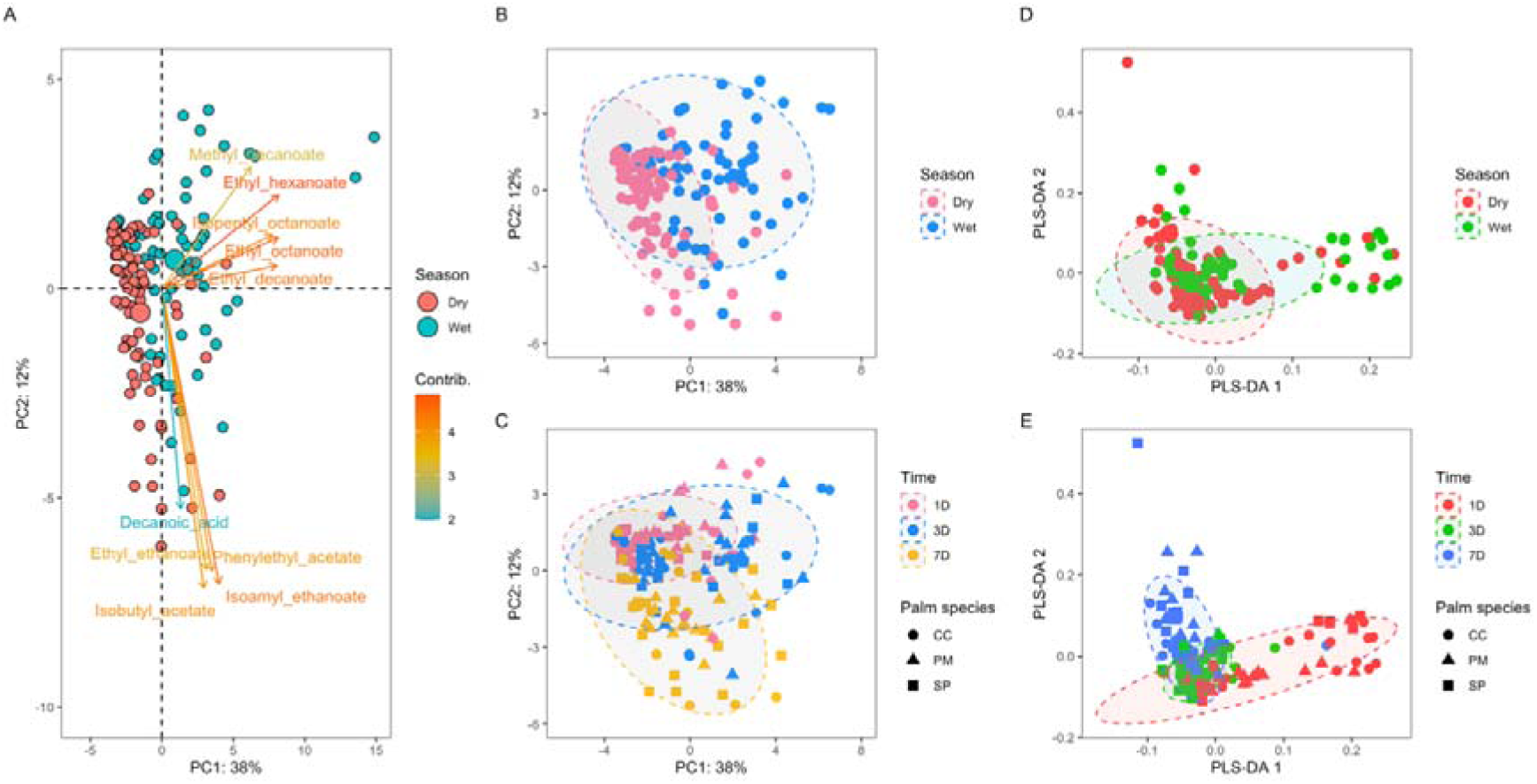
Profile of volatile compounds and small molecules of palm wine showing the impact of season, palm species, and fermentation time to their formation. A. PCA biplot shows the correlation of ecological factors to specific compounds; B. Profile of volatile compounds was significantly clustered across season; C. The volatile compound composition was clearly differentiated by the fermentation time; D-E. Small molecule composition was clustered by season and fermentation time.

We measured small molecules to obtain wider overview of aroma and flavour construction in palm sap fermentation. Proton NMR was conducted to predict the wider small molecules by clustering the proton peaks across all variables (**Fig. 4D-E**). We obtained 181 bins of peaks that were clustered into 59 groups of compounds (buckets), listed in **Supplementary Table S3**. The peak intensity occurred in the ranges of 1-1.5 ppm, 2-2.5 ppm, 3.2-4.4 ppm, and 4.6-5.6 ppm, indicating the major molecules of palm wine, such as ethanol, sugars, amino acids, and organic acids. Partial least discriminant analysis (PLSDA) provided a significant separation through class that can easily show the effect of factors. After discriminating bins of all samples, the general pattern reflected the profile given by the volatile compound profiles, showing a distinct cluster in all variables. Season of collection clearly grouped samples with model accuracy (cross-validation) at 0.863, palm tree species separated into three clusters with model accuracy at 0.826, while separation in fermentation time showed accuracy at 0.851. These accuracies confirm that the separation model was valid and could accurately classify small molecules in palm wine across sample parameters.

To understand aroma and flavour formation affected by our classified variables, three-way ANOVA analysis was used to compare the change of compound composition and intensity (**Fig 5**). We did not include the data from small molecules of 1H’NMR data due to the difficulty in calculating the absolute concentration by this method. The intensities of esters, benzenes, and aldehydes were significantly different between seasons and fermentation time. Ethyl decanoate and ethyl octanoate were the highest concentration and higher in the wet season and tend to decrease at the final fermentation timepoint (**Fig. 5A**). A Comparison between palm species and the change of season showed a significant influence in esters, which increased at the end of fermentation (**Fig. 5B**). Groups of compounds were related to the alteration all variables where ethyl octanoate was the most obvious volatile compound showing in its intensity (**Fig. 5C**).

**Fig 5.**
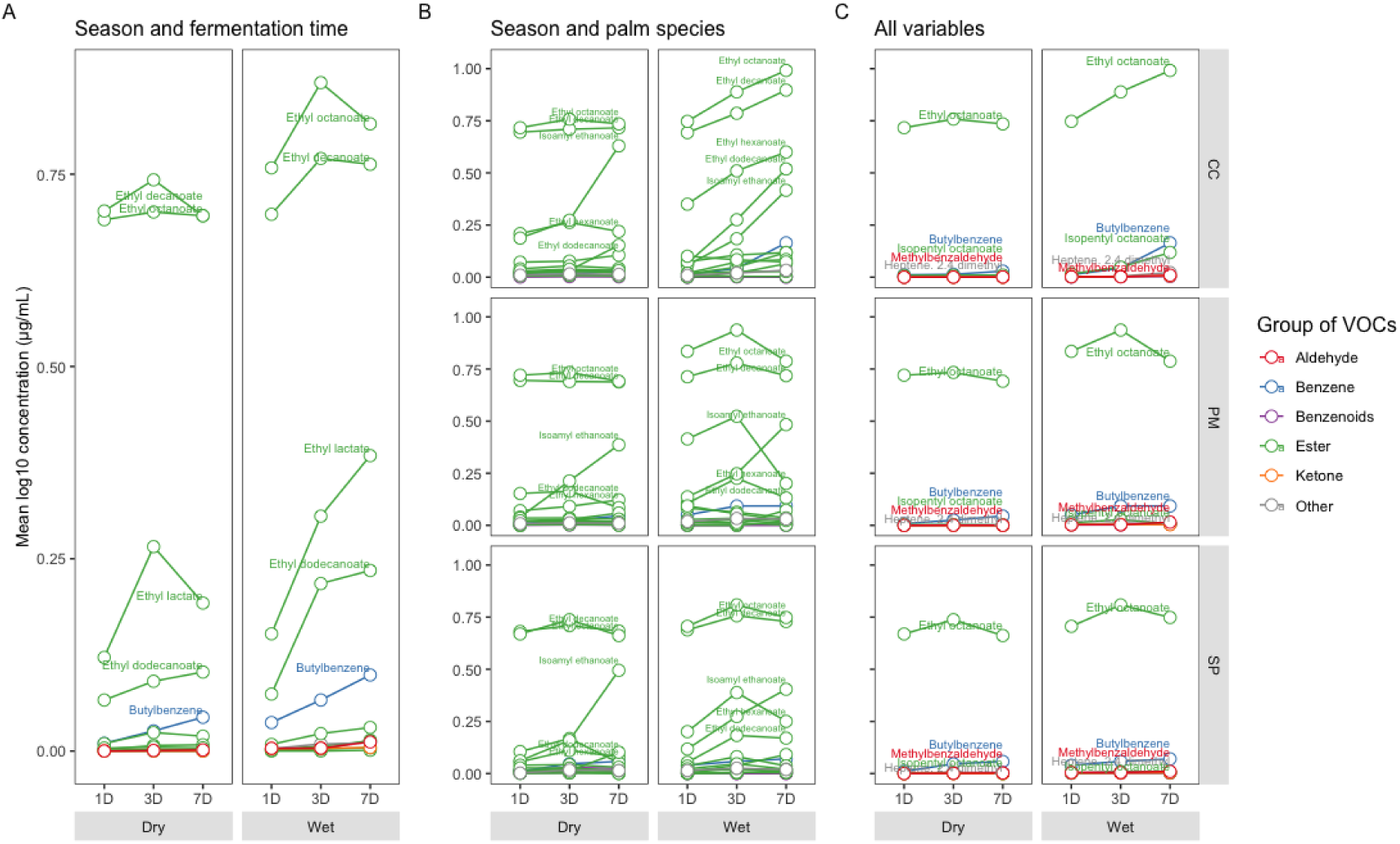
The top 10 significant volatile compounds across season, fermentation time, and palm species measured by GC-MS and selected by ANOVA. A) Esters, benzenes and aldehydes were significantly related to the change of season and fermentation time based on ANOVA-two way; B) Across palm species and the change of season, the highest effect was observed upon ester; C) In three-way ANOVA, ethyl octanoate was the most obvious compound that was highly affected by all variables.

### 3.5 Spoilage species have negative impacts on ester formation

The relationship of microbial communities and metabolites to season of collection, palm species and time of fermentation was demonstrated, yet our understanding of the contribution of microbial communities on aroma and flavour formation was incomplete. Through the correlation analysis, we simulated the links between the microbial communities and compound composition to understand the factors influencing their importance to palm sap fermentation (**Fig. 6**). In term of core microbial taxa, a positive correlation was shown between bacterial and fungal taxa (**Fig. 6A**). When compared to the top volatile compounds, the genera of *Zymomonas* had the highest contribution to the top volatile compounds, while reads for *Lentilactobacillus*, *Lactobacillus, Liquorilactobacillus*, and *Acetobacter* genera have a positive correlation on isoamyl acetate and ethyl ethanoate (**Fig. 6B**). All fungal core taxa were observed to contribute to the top volatile compounds with *Saccharomyces* spp. contributing the most (**Fig. 6C**). We further analysed the correlation of microbial taxa and metabolites, that were significantly affected by season, suggested by previous three-way ANOVA and DESeq2 results (**Fig. 6D-E**). The presence of *Leuconostoc-, Zymobacter-,* and *Serratia*-genus associated reads negatively affected esters while *Lactococcus* has significant effect to non-ester, isobutyl acetate and isoamyl ethanoate. In fungal communities, the human-related species *Candida tropicalis*, were negatively correlated with the presence of esters and the cosmopolitan yeast *Blastobotyris*, had a negative correlation to isobutyl acetate and isoamyl ethanoate. These taxa are likely related to the change of the aroma and flavour formation of palm wine during the seasonal sap tapping and may act as spoilage species, downgrading the quality of palm wine characteristics.

**Fig. 6.**
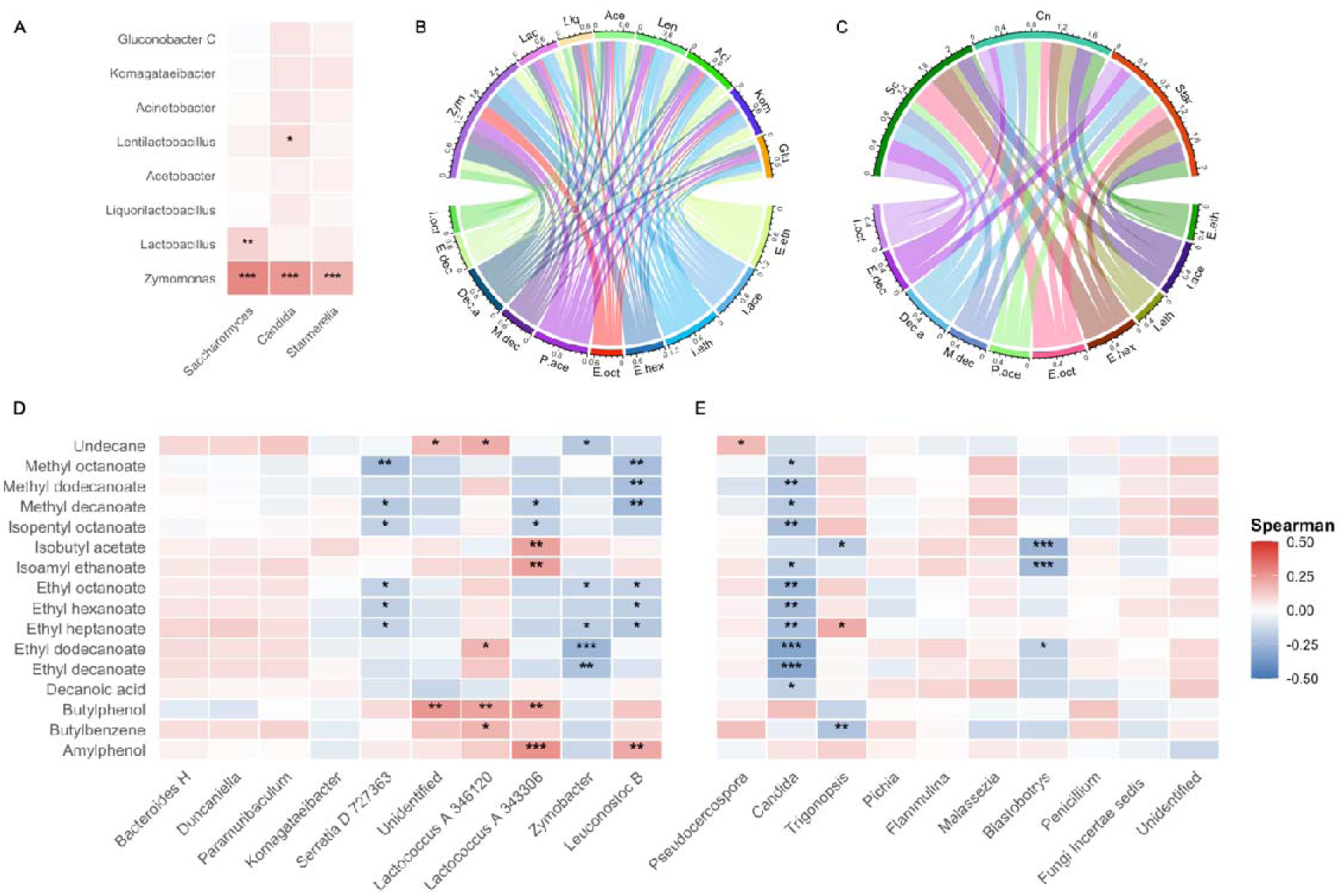
Correlation analysis of microbial communities and volatile compounds in palm wine. A) Correlation heatmap of core species of bacteria and fungi; B-C) Cord diagram showing the correlation of core ASVs (Bacteria [B]: Zym = *Zymomonas*, Lac = *Lactobacillus*, Liq = *Liquorilactobacillus*, Ace = *Acetobacter*, Len = *Lentilactobacillus*, Aci = *Acinetobacter*, Kom = *Komagataeibacter*, Glu = *Gluconobacter*; Fungi [C]: Sc = *Saccharomyces*, Cn = *Candida*, Star = *Starmerella*) and top 10 of the highest concentration of VOCs (I.oct = Isoamyl octanoate, E.dec = Ethyl decanoate, Dec.a = Decanoic acid, M.dec = Methyl decanoate, E.oct = Ethyl octanoate, E.hex = Ethyl hexanoate, I.eth = Isoamyl ethanoate, I.ace = Isoamyl acetate, E.eth = Ethyl ethanoate); D-E) Correlation heatmap of the most significant VOCs in season and palm species retrieved from three-way ANOVA analysis, and the top 10 significant genera of microbial communities (*p*-value= 0.05) from DESeq2 analysis.

## 4. Discussion

Palm sap fermentation is a rapid and spontaneously initiated activity driven by natural microbial communities that are sensitive to environmental and ecological factors. Our analysis of palm sap fermentation in Bali has uncovered multiple microbial dynamics in a unique and culturally important food production system. In this study, we elucidated the influence of tree species and palm sap collection season on microbial composition and associated these changes with the metabolite profile during the palm sap fermentation. Our findings suggest that seasonality strongly shapes the microbial structure of palm sap fermentation, leading to a loss of diversity indices observed from the dry to wet season. The volume and composition of fresh palm sap are influenced by seasonal changes (Ziadi et al., 2014) and in this study, the concentration of sugar and titratable acidity differentiated sap by the season collected. This alteration impacts the fungi and bacteria able to grow in palm sap.

Palm trees are a perennial plant that grow throughout the year, and experience two seasons (wet and dry) in tropical regions. The wet season is generally higher temperature and has higher available water and humidity. The dry season by contrast, has warm temperatures, but with more extensive light exposure. In other plants, seasonal factors can mediate microbial communities of plant spatial ecosystem from soil to the aerial parts of the plant (Ding & Melcher, 2016; He et al., 2024; Waldrop & Firestone, 2006). Soil microbial communities are sensitive to soil conditions (Fiore-Donno et al., 2024; Thoms & Gleixner, 2013; Waldrop & Firestone, 2006) and spatial microbes, such as the endophytic and phyllosphere microbiota, are also vulnerable (Classen et al., 2015; He et al., 2024). Abiotic stress can also affect plant-insect relationships (Pineda et al., 2013). Insects play an important role in the dispersal of microbial communities on the surface of palm trees, and palm leaves or fronts are often added to fresh palm sap as the fermentation takes place (Sumerta et al., 2025). Together, these seasonal factors provide perspectives of microbial dynamics based on seasonal collection and fermentation of palm sap and likely affects the complexity of palm wine characteristics.

In many fermentation systems, microbial dominance plays a critical role in shaping microbial dynamics, directing the fermentation products. Core species such as *S. cerevisiae* and *Z. mobilis* are dominant throughout palm sap fermentation process. *Saccharomyces* yeasts have different mechanisms to dominate various fermentative environments through physical exclusion, genetic characteristics, and strong metabolic pathway in mixed culture environments (Albergaria & Arneborg, 2016; Pérez-Torrado et al., 2017), while *Z. mobilis* is a facultative anaerobic bacterium that has high tolerance to ethanol, sugar content, and phenolic acids (Banta et al., 2020; Gu et al., 2015; Rutkis et al., 2022). In some cases, *Z. mobilis* can outperform the growth and ethanol yield of *S. cerevisiae* (Banta et al., 2020). Microbial succession is evident during fermentation, giving temporal variations in palm sap fermentation (**Fig. 3E-F**). Microbial species occur and vary as a result of environmental conditions, showing a co-occurrence pattern that assembles species to interact and respond (Fuhrman, 2009). In palm wine, we have shown a shift in the microbial communities associated with changes in the composition of metabolites and other small molecules. Although the dry season had lower microbial richness indices than the wet season, we observed altered formation of aroma and flavours, especially ester production.

Esters are important for flavour of fermented beverages. We show here that esters were altered by all variables studied, but their fluctuation was not significantly related to the core taxa excepting to *Candida* spp., which was negatively correlated with esters. *Candida* spp. occurrence was higher in the dry season, reduced moisture and related changes favour its growth. The presence of spoilage yeast can change the aromatic profile as certain non-*Saccharomyces* express esterase activity that hydrolyse volatile esters (Belda et al., 2017; Fleet, 2011). Esters are frequently produced by *S. cerevisiae* due to its *ATF1* gene encoded enzymes that are responsible for synthesising volatile acetate esters, facilitating the dispersal of this yeast cells through insect movements (Christiaens et al., 2014; Verstrepen et al., 2003). Over all ester constituents, ethyl octanoate was consistently significant across all variables and comparison. This compound has a floral aroma and is commonly used as a flavour enhancer and fragrance ingredient and is found in spirits, wines, and fruits (J. Liu et al., 2021). This prominent volatile organic compound is suggested as a biomarker to quantify the impact of seasonal variability, on the aroma and flavour of fermented palm sap and associated products.

We show here that seasonal factors strongly affect the fermentation process of palm wine and thus contribute to perspectives on managing microbial fermentation in plant-substrates. Season and palm species are the main factors that influence differentiation of microbial and metabolite composition of palm sap while fermentation time is likely less significant to the change of dominant species in palm sap fermentation. These variables facilitate a dynamic environment within the local geographic area, leading to microbial responses and specific metabolite production, while in a bigger picture this change can capture product specificity based on geographic origin (Bokulich et al., 2016). Microbial species exhibit spatial and temporal dynamics that can influence how sampling strategies are designed to capture ecological fluctuations and to manage the stability of the fermentation product quality (Barata et al., 2012), and we show how this occurs in palm sap fermentation.

## 5. Conclusion

Season of sap collection significantly shapes the microbial community and metabolite formation in palm wine, directly influencing the differentiation of the product aroma and flavour profile. Most notably, microbial transformations drive the production of flavoursome esters with ethyl octanoate emerging as a key volatile biomarker for these seasonal variations. Furthermore, the complex and dynamic microbial and metabolic formation is inherently linked to the specific palm species and the dominant taxa ultimately dictate the microbial diversity in palm sap fermentation.

## Data availability

The amplicon datasets of bacteria and fungi generated for this study are available on the NCBI Bioproject PJRNA1480356 and PRJNA1483710, respectively.

## Acknowledgements

The authors would like to acknowledge Melbourne High Performance Computing for enabling microbiome analysis, and an Australia Award for funding and supporting INS. We would like to thank David Keizer and Sunnia Rajput for helping with NMR data acquisition at Bio21, Pangzhen Zhang and Xavier Luo for advice on HS-GC/MS data analysis. BRIN colleagues, Yeni Yuliani, who helped sample export and Prof. I Made Sudianan – Research Center for Applied Microbiology (RCAM) for facilitating material transfer agreement of the exported samples to Melbourne. The farmers in Bali are gratefully acknowledged for their time, expertise in collection of samples and enthusiasm for the project.

---

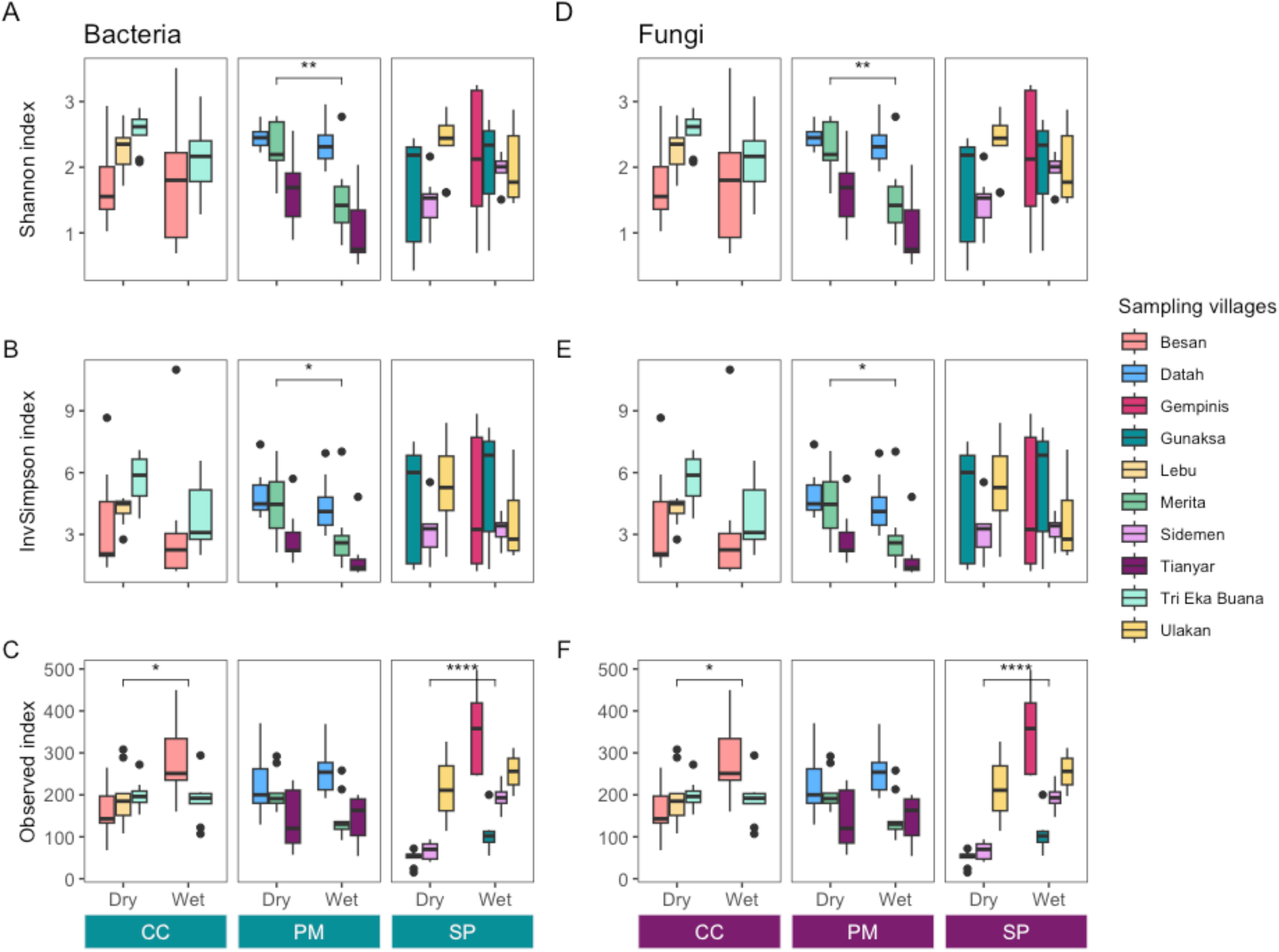

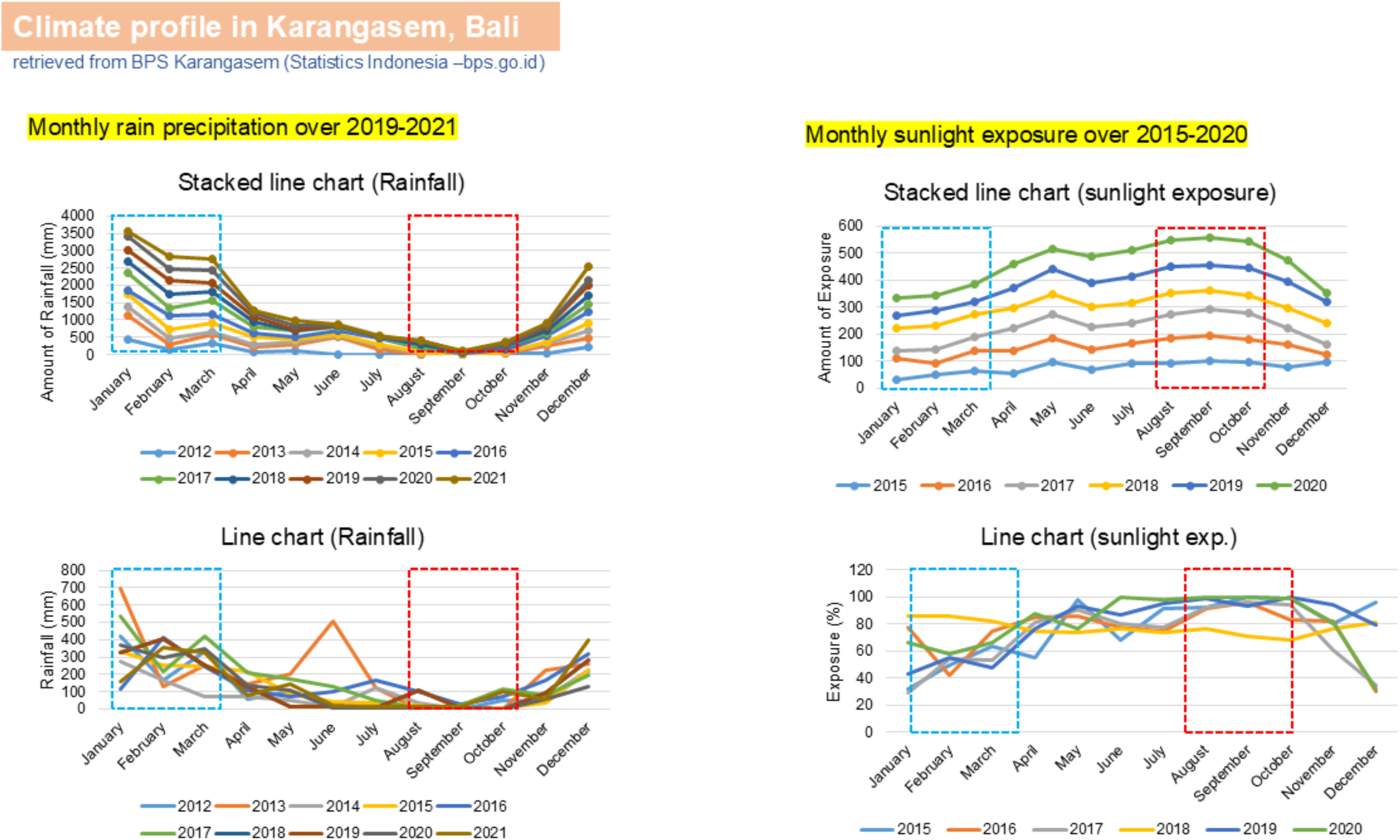

## Notes

### Competing Interest Statement

The authors have declared no competing interest.

